# A bacterial assay for rapid screening of IAA catabolic enzymes

**DOI:** 10.1101/681379

**Authors:** Brunoni Federica, Collani Silvio, Šimura Jan, Schmid Markus, Bellini Catherine, Ljung Karin

**Author notes:** These authors contributed equally to this work (Brunoni Federica, Collani Silvio). These authors contributed equally to this work (Bellini Catherine, Ljung Karin).

## Abstract

**Background:** Plants rely on concentration gradients of the native auxin, indole-3-acetic acid (IAA), to modulate plant growth and development. Both metabolic and transport processes participate in the dynamic regulation of IAA homeostasis. Free IAA levels can be reduced by inactivation mechanisms, such as conjugation and degradation. IAA can be conjugated via ester linkage to glucose, or via amide linkage to amino acids, and degraded via oxidation. Members of the UDP glucosyl transferase (UGT) family catalyze the conversion of IAA to indole-3-acetyl-1-glucosyl ester (IAGlc); by contrast, IAA is irreversibly converted to indole-3-acetyl-L-aspartic acid (IAAsp) and indole-3-acetyl glutamic acid (IAGlu) by Group II of the GRETCHEN HAGEN3 (GH3) family of acyl amido synthetases. DIOXYGENASE OF AUXIN OXIDATION (DAO) irreversibly oxidizes IAA to oxindole-3-acetic acid (oxIAA) and, in turn, oxIAA can be further glucosylated to oxindole-3-acetyl-1-glucosyl ester (oxIAGlc) by UGTs. These metabolic pathways have been identified based on mutant analyses, *in vitro* activity measurements, and *in planta* feeding assays. *In vitro* assays for studying protein activity are based on expressing Arabidopsis enzymes in a recombinant form in bacteria or yeast followed by recombinant protein purification. However, the need to extract and purify the recombinant proteins represents a major obstacle when performing *in vitro* assays.

**Results:** In this work we report a rapid, reproducible and cheap method to screen the enzymatic activity of recombinant proteins that are known to inactivate IAA. The enzymatic reactions are carried out directly in bacteria that express the recombinant protein. The enzymatic products can be measured by direct injection of a small supernatant fraction from the bacterial culture on ultrahigh-performance liquid chromatography coupled to electrospray ionization tandem spectrometry (UHPLC-ESI-MS/MS). Experimental procedures were optimized for testing the activity of different classes of IAA-modifying enzymes without the need to purify recombinant protein.

**Conclusions:** This new method represents an alternative to existing *in vitro* assays. It can be applied to the analysis of IAA metabolites that are produced upon supplementation of substrate to engineered bacterial cultures and can be used for a rapid screening of orthologous candidate genes from non-model species.

## Background

The phytohormone auxin is a master regulator of plant growth and development affecting a plethora of different processes during a plant’s life-cycle. Auxin acts in a concentration-dependent manner and its maximum and minimum requirements vary among different plant tissues [1] and specific cell types [2-4]. Thus, the optimum endogenous auxin level must be tightly and finely controlled. Several regulatory mechanisms operate in plant cells to modulate the levels of the bioactive form of this hormone [5].

The major active auxin found in plants is indole-3-acetic acid (IAA). IAA is synthesized in plants primarily by the tryptophan (Trp)-dependent pathway, although evidence for a Trp-independent mechanism of IAA biosynthesis has also been reported [5,6]. Apart from *de novo* synthesis, inactivation of the hormone also participates in the dynamic regulation of auxin metabolism, increasing the complexity of auxin homeostasis in cells. The two main routes of auxin inactivation are the conjugation of IAA to amino acids and sugars, and the oxidation of IAA to oxindole-3-acetic acid (oxIAA) [7,8]. IAA conjugates are generally considered to be storage reserves of auxin as some of them can be hydrolyzed back to release free IAA [5]. IAA can be converted to amide conjugates by conjugation with amino acids by Group II members of the GRETCHEN HAGEN 3 (GH3) family of IAA-amide synthases [9,10]. However, IAA conjugated with both aspartate (indole-3-acetyl-L-aspartic acid, IAAsp) and glutamate (indole-3-acetyl-L-Glutamic acid, IAGlu) are non-hydrolysable and their formation leads to an irreversible removal of IAA from the active pool [11,12]. In addition, IAA can be reversibly conjugated via ester linkages to glucose by UDP-glucosyl transferases to produce indole-3-acetyl-1-glucosyl ester (IAGlc), where the *Arabidopsis thaliana* enzymes UGT84B1 and UGT74D1 are the strongest candidates [13-15]. The oxidative pathway is the major IAA catabolic pathway in Arabidopsis [12,16,17] and the irreversible formation of oxIAA was only recently ascribed to the activity of members of the 2-oxoglutarate-dependent-Fe(II) dioxygenases, DIOXYGENASE FOR AUXIN OXIDATION (DAO) [18,19]. oxIAA can be further conjugated to glucose by UGT74D1 in Arabidopsis, to produce oxindole-3-acetyl-1-glucosyl ester (oxIAGlc) [15].

The identification of the different metabolic pathways has mainly been based on mutant analysis, metabolic profiling, *in planta* feeding assays, and measuring of enzyme activities *in vitro*. Metabolite feeding experiments have been carried out by supplying labeled IAA followed by mass spectrometry (MS) to identify *de novo* synthesized IAA catabolites and conjugates [12,16,18,20,21]. When Arabidopsis seedlings were fed with low concentrations of labeled auxin, IAA was mainly degraded to oxIAA and oxIAGlc, while higher levels of labeled IAA induced conjugation to IAAsp and IAGlu [12]. Similar metabolic analysis of *dao1* loss-of-function mutant revealed that the reduction of IAA oxidation in this mutant did not change the levels of free IAA, whereas the IAA conjugates, IAAsp and IAGlu, accumulated at much higher levels in the *dao1* mutant than in wild type seedlings [18]. These findings provide evidence to support the idea that DAO- and GH3-mediated catabolic pathways influence each other’s activities to maintain auxin homeostasis [22]. The functional characterization of GH3, UGT and DAO proteins *in planta* can be challenging due to large families and functional redundancy. Therefore, *in vitro* assays have previously been conducted with recombinant proteins in order to elucidate the molecular function of GH3, UGT and DAO enzymes during IAA inactivation [9,13-15,19,23]. GH3 enzymatic activity with various amino acid substrates has been examined using High Performance Liquid Chromatography (HPLC)- and thin-layer chromatography (TLC)-based assay methods in Arabidopsis [9] and in other plant species [24,25]. Results from these studies have demonstrated that each GH3 enzyme has slightly different substrate specificity and probably contributes to the formation of several types of IAA-amino acid conjugates. Nonetheless, *in vitro* assays have revealed IAAsp to be a major conjugate formed by GH3.6, and that GH3.17 appears to favor glutamate over aspartate [9]. More recently, the development of a LC-MS-based assay allowed a steady-state kinetic analysis of several rice GH3 enzymes [26-28]. Several studies have shown that recombinant Arabidopsis UGT84B1 and UGT74D1 proteins catalyze the glucosylation of oxIAA and IAA to their corresponding glucosides *in vitro* [13-15,23]. Moreover, recombinant UGT74D1 protein demonstrated to have a strong glucosylation activity towards various auxin-related compounds, but its activity is stronger with oxIAA than with IAA as a substrate [15].

IAA conjugates and oxIAA are also predominant IAA metabolites, not only in other *Viridiplantae*, such as bryophytes [29] and algae, but also in cyanobacteria [30], indicating that the mechanisms of IAA inactivation occurred very early during evolution [8].

The *in vitro* enzyme assays are usually based on heterologous expression of plant genes in bacteria or yeast, followed by recombinant protein purification. This procedure can be very time-consuming and needs specific equipment and reagents to purify proteins on a large scale. Furthermore, the solubility and stability of the protein of interest can be problematic and lead to either poor yield or inactivation of the protein.

Here we introduce a reliable method for rapidly screening the activity of IAA conjugation and oxidation enzymes where the reactions have been performed directly in bacteria using recombinant proteins (Fig. 1a). This method allows the detection and quantification of enzyme products without any purification step, after direct injection of a small supernatant fraction of the bacterial liquid culture, using ultrahigh-performance chromatography coupled to electrospray ionization tandem spectrometry (UHPLC-ESI-MS/MS). Our analysis shows that the products of interest were more abundant in the supernatant than in the pellet. The combination of these methods creates a faster, more reproducible and cheaper alternative to detect IAA catabolic products compared to the methods adopted so far. The proposed method provides a powerful alternative to screening orthologous candidate genes from non-model species, which is an important challenge due to the exponential increase of the sequenced genomes.

**Fig. 1.**
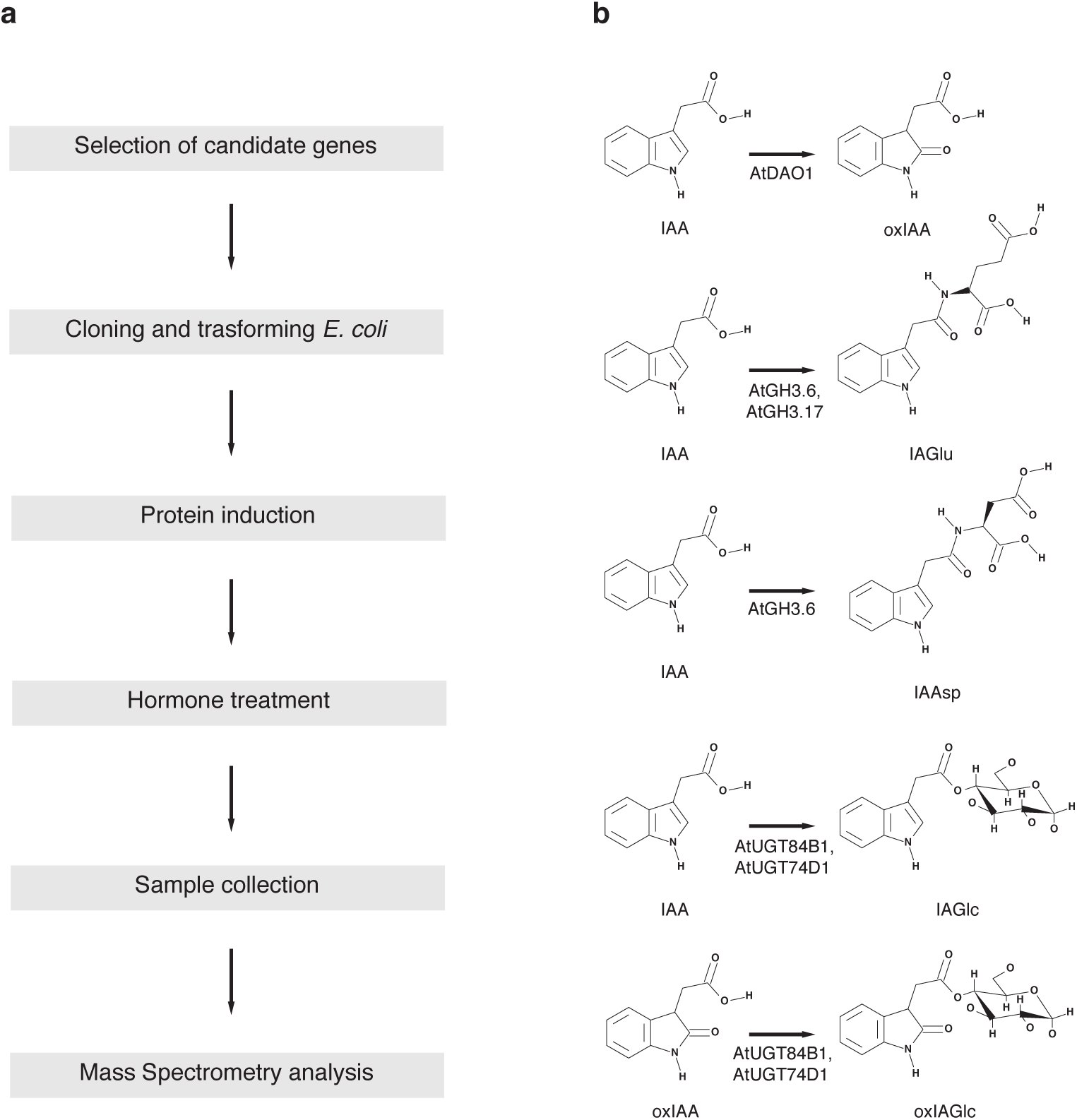
**a** Schematic representation of the bacterial enzymatic assay. **b** Enzymes and biosynthetic pathways that inactivate IAA and that are studied in this article.

## Methods

### Cloning and protein expression of AtDAO1, AtGH3.6, AtGH3.17, AtUGT84B1 and AtUGT74D1

Full-length cDNAs of *AtDAO1* (At1g14130), *AtGH3.6* (At5g54510), *AtGH3.17* (At1g28130), *AtUGT84B1* (At2g23260) and *AtUGT74D1* (At2g31750) were amplified with gene-specific primers containing additional *XbaI* and *HindIII* and *NheI* sites (<u>Additional File 1: Table S1</u>) using Arabidopsis cDNA as template. The cDNA fragments were cloned into the *XbaI* and *HindIII* or *NheI* restriction sites of a pETM-11 vector to generate recombinant fusion proteins with an N-terminal 6x His tag. The constructs were transformed into *Escherichia coli* BL21 (DE3). An *E. coli* strain carrying the pETM-11 vector containing the GFP coding sequence was used as control. For protein expression, *E. coli* containing the plasmids were grown in shaker flasks (250 rpm at 37°C) containing Luria Broth supplemented with 50 μg/ml (w/v) kanamycin to OD_600_ between 0.4 and 0.6. The production of recombinant proteins was induced by the addition of 0.1 mM isopropyl-β-D-thiogalactopyranoside (IPTG). To monitor the correct protein induction, cells were grown for 6 hours after IPTG induction at 20°C with constant shaking at 250 rpm and bacterial cultures were harvested for SDS/PAGE and Western blot analysis.

### SDS/PAGE and Western blot analysis

To verify the induction of recombinant proteins, 2 ml of bacterial culture were harvested before and 6 hours after IPTG induction; cells were pelleted by centrifugation at 4000 rpm for 5 minutes. The pellet was washed and resuspended with 1 ml 1x PBS buffer (1 mM KH_2_PO_4_, 10 mM NaHPO_4_, 137 mM NaCl, 2.7 mM KCl) and cells lysed by sonication. Cell debris was pelleted by centrifugation (14000 rpm, 30 minutes, 4°C) and the supernatant was collected. Ten μl of 4x Laemmli sample buffer (Biorad) was added to 30 μl of supernatant and denatured by boiling at 95°C for 5 minutes. Samples were then loaded on a 12% (w/v) SDS/PAGE gel and, after gel run, transferred to a PVDF membrane by wet electroblotting. For detection of recombinant proteins, the PVDF membrane was blocked in 1x TBS-T with 5% skimmed milk for 2 hours at room temperature. After blocking, the membrane was incubated with anti-6x His antibody horseradish peroxidase (HRP) conjugate (Sigma A7058-1VL, 1:10000 dilution in 1x TBS-T with 5% skimmed milk) for 2 hours at room temperature and washed for 10 minutes, four times, in 1x TBS-T. Signals were detected by an enhanced chemiluminescent (ECL) detection system (GE Healthcare RPN2232). ECL signals were captured by a charge-coupled device (CCD) camera (Azure c600) (<u>Additional File 2: Figure S1</u>).

### Enzyme assay

To test the enzymatic activity of recombinant proteins, over-night induced bacterial cultures were supplemented with substrate in the culture medium. For AtDAO1, a culture of AtDAO1-expressing bacteria was supplemented with 0, 0.001, 0.01, or 0.1 mM IAA and with or without DAO cofactor mixture [5 mM 2-oxoglutarate, 5 mM ascorbate, 0.5 mM (NH_4_)_2_Fe(SO_4_)_2_]. For the first trial, 1 ml samples were taken before (at time 0, T0) and after over-night incubation at 20°C with or without 0.001 mM IAA and with or without DAO cofactor mixture, centrifuged to pellet (4000 rpm for 5 minutes) and the resulting supernatant and pellet fractions were stored at −20°C and analyzed for oxIAA production by UHPLC-ESI-MS/MS as described below. The following experiments were carried out by analyzing only the supernatant fraction. Subsequently, the AtDAO1-expressing bacterial culture was treated with or without 0.01 mM IAA, with or without the DAO cofactor mixture and kept at 20°C or 30°C. Samples were taken at 1 and 6 hours after incubation. Samples from GFP-expressing bacterial cultures and media without bacteria were used as negative controls. For AtGH3s, cultures of AtGH3.6- or AtGH3.17-expressing bacteria were kept at 20°C and treated with or without 0.1 mM IAA and with GH3 cofactor mixture (1 mM Glutamic acid, 1 mM Aspartic acid, 3 mM ATP, 3 mM MgCl_2_). Samples from GFP-expressing bacterial cultures were used as negative controls. Samples were collected 6 hours after incubation. For AtUGTs, cultures of AtUGT84B1- or AtUGT74D1-expressing bacteria were kept at 20°C and treated with or without 0.1 mM IAA and/or oxIAA and with UGT cofactor mixture (1 mM UDP-glucose, 2.5 mM MgSO_4_, 10 mM KCl). Samples were collected 6 hours after incubation. Samples from GFP-expressing bacterial cultures were used as negative controls. All the experiments were carried out with shaking at 250 rpm in the dark and repeated in triplicate.

### UHPLC-ESI-MS/MS analysis

Supernatant fractions from bacterial cultures were analyzed according to the UHPLC-ESI-MS/MS method for auxin metabolic profiling described in [17,31], with modifications. The fractions were mixed using a vortex, and 2 μl per sample were transferred in insert-equipped vials and spiked with 5 μl of a 13C_6_ labeled internal standard mixture, containing 5 pmol of labeled IAA, oxIAA, IAAsp, IAGlu, IAGlc and oxIAGlc. Samples were further diluted in 20% (v/v) methanol up to 40 μl and 2 μl per sample were injected onto a reversed-phase column (Kinetex C18 100A, length 50 mm, diameter 2.1 mm, particle size 1.7 μm; Phenomenex) and eluted using a 14.5 min gradient comprising 0.1% acetic acid in water (A) and 0.1% acetic acid in methanol (B) at a flow rate of 0.25 ml/min; column temperature was set to 30°C. The following binary linear gradient was used: 0 min, 10:90 A:B; 10.0 min, 50:50 A:B; during 11-12.2 min the column was washed with 2:98 A:B; and between 12.5-14.5 min re-equilibrated to initial conditions, 10:90 A:B. Quantification was obtained by multiple reaction monitoring (MRM) of the precursor ([M+H]+) and the appropriate product ion (<u>Additional File 3: Table S2</u>). Mass spectrometry analysis and quantification were performed by an LC-MS/MS system comprising a 1290 Infinity Binary LC System coupled to a 6490 Triple Quad LC/MS System with Jet Stream and Dual Ion Funnel technologies (Agilent Technologies, Santa Clara, CA, USA).

## Results and discussion

### Optimization and validation of bacterial assay with AtDAO1

To optimize and validate the proposed method, we used *E. coli* cultures expressing recombinant AtDAO1 to test the effect of different experimental conditions on oxIAA production. AtDAO1 catalyzes the oxidation of IAA (Fig. 1b) and this reaction represents the most important mechanism for IAA inactivation in plants [18,21,29,30]. Recombinant AtDAO1 was expressed in bacterial cultures as an N-terminally tagged hexahistidine fusion protein and its expression was confirmed by Western blot analysis (<u>Additional File 2: Figure S1</u>). In the first experiment, the enzymatic activity of recombinant AtDAO1 was tested by supplementing bacterial cultures expressing AtDAO1 with 0.001 mM IAA and measuring the levels of oxIAA in the supernatant and in the pellet fractions separately by UHPLC-ESI-MS/MS. Previous work has shown that the *in vitro* catalytic activity of recombinant DAO proteins [19,32] requires, in addition to the substrate IAA, the co-substrate 2-oxoglutarate, and the cofactors (NH_4_)_2_Fe(SO_4_)_2_ and ascorbate in the reaction mixture. We therefore carried out, the assay by adding IAA to the bacterial cultures with or without a mixture containing co-substrate and cofactors (DAO cofactor mixture). Bacterial cultures expressing the GFP protein supplemented with IAA in the presence or absence of the DAO cofactor mixture were used as negative controls. Untreated bacterial cultures were used as mock samples. Bacterial cultures were sampled prior to the treatment with IAA (T0) and after over-night incubation at 20°C with the substrate, and with or without the DAO cofactor mixture. Under these conditions, the supernatant fraction exhibited a strong accumulation of oxIAA in all the samples compared with the pellet (Fig. 2a). Our analysis shows the concentration of oxIAA to be higher in the culture media of bacterial cells transformed with *AtDAO1* that were simultaneously incubated with IAA and DAO cofactor mixture than in the samples taken from the same bacterial culture that was treated with IAA alone (Fig. 2a). However, background levels of oxIAA were observed in the culture media of AtDAO1-expressing bacteria that were not treated with IAA (mock), prior treatment (T0) and in all GFP samples (Fig. 2a). Low levels of IAA were detected in culture media before treatment with exogenous IAA (<u>Additional File 4: Figure S2</u>). Moreover, the basal accumulation of oxIAA that we observed in bacterial cultures could have been derived from the spontaneous non-enzymatic conversion of IAA into oxIAA, as has previously been reported in control samples from *in vitro* AtDAO assays [19]. Our results showed that AtDAO1 activity could be monitored by direct analysis of the levels of oxIAA that accumulate in the supernatant fraction of the AtDAO1-expressing bacterial cultures that had been supplemented simultaneously with low levels of exogenous IAA and DAO cofactor mixture. Therefore, in all further tests only the supernatant fractions were analyzed.

**Fig. 2.**
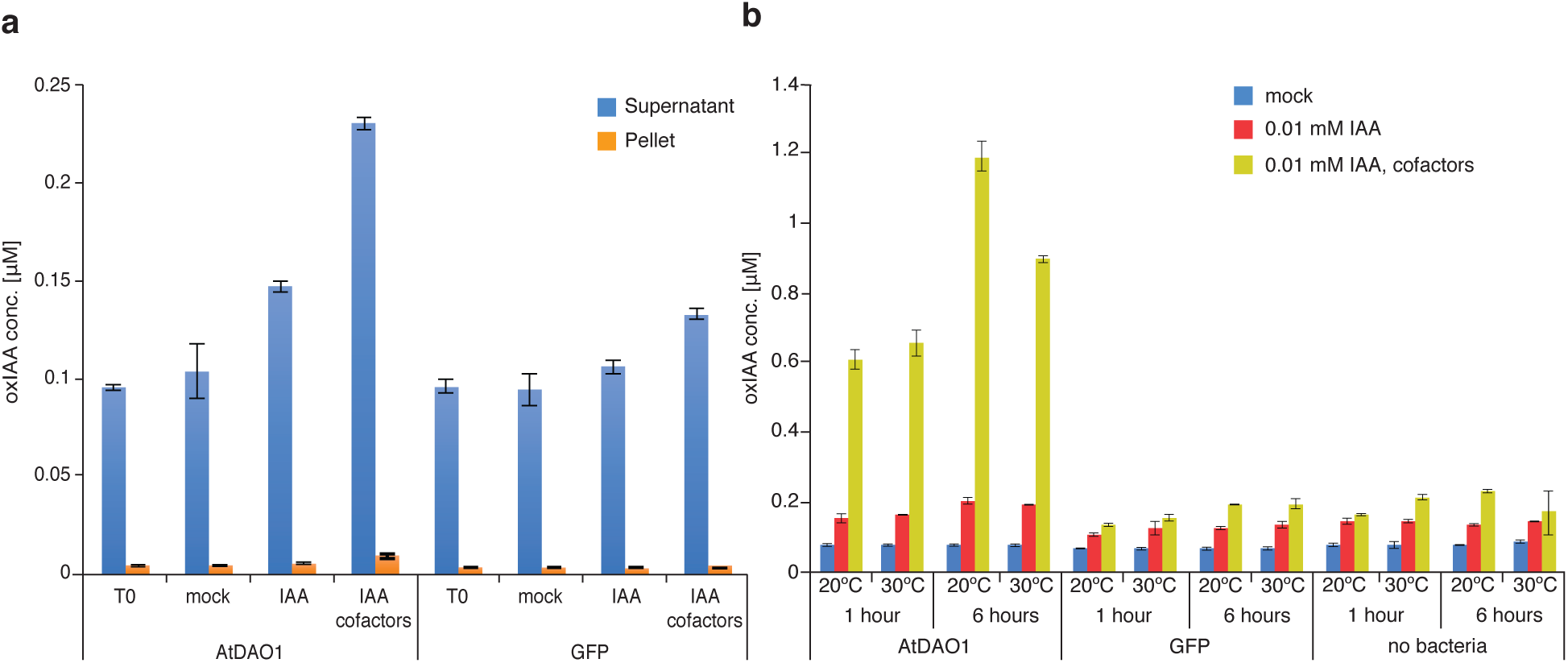
Bacterial assay of IAA oxidation mediated by recombinant AtDAO1. **a** oxIAA concentrations in supernatant or pellet fractions from AtDAO1- or GFP-expressing bacterial cultures supplemented with 0.001 mM IAA and with or without the DAO cofactor mixture. T0: samples taken before supplementation with exogenous IAA. **b** oxIAA concentrations in supernatant fractions from AtDAO1- or GFP-expressing bacterial cultures supplemented with 0.01 mM IAA and with or without the DAO cofactor mixture for 1 or 6 hours and incubated at either 20°C or 30°C. Bacterial cultures without IAA and the DAO cofactor mixture were used as mock samples. ‘No bacteria’, are media in which bacteria were not cultured. Mean ± SD (*n*=3).

In the following experiment, we investigated the effect of temperature and exposure time to exogenous IAA and the DAO cofactor mixture on AtDAO1-mediated oxIAA production. In previous studies, *in vitro* DAO assays were performed by incubating the reaction mixture containing the purified recombinant DAOs at 30°C for 1 hour [19,32]. We therefore carried out an experiment in which AtDAO1- and GFP-expressing bacterial cultures were incubated at 20°C or 30°C with IAA and in the presence or absence of the DAO cofactor mixture. Samples were taken after 1 or 6 hours of treatment with IAA. To reduce the impact of the natural non-enzymatic oxidation of IAA on the overall oxIAA production, bacterial cultures were treated with a higher concentration of IAA (0.01 mM) than that adopted in the initial experiment (0.001 mM). Untreated bacterial cultures were used as mock samples and culture media that had not been inoculated with bacteria (designed ‘no bacteria’) were sampled and analyzed as an additional control to monitor the background oxIAA formation that derives from the naturally occurring oxidation of IAA. We observed a very strong accumulation of oxIAA in AtDAO1 samples taken from cultures that were treated with IAA and the DAO cofactor mixture, compared with AtDAO1 from samples treated with IAA only (Fig. 2b), indicating that the simultaneous supplementation of the DAO cofactor mixture and a higher concentration of IAA significantly enhanced AtDAO1-mediated oxIAA production. This finding was also confirmed when AtDAO1-expressing bacterial cultures were incubated with 0.1 mM IAA (Fig. 3a). The concentration of oxIAA was higher in AtDAO1 samples after prolonged treatment (6 hours) with substrate and the DAO cofactor mixture, than in samples that had a shorter exposure time (1 hour) (Fig. 2b). Furthermore, oxIAA levels were slightly higher in the samples that were incubated at 20°C compared with samples from cultures kept at 30°C (Fig. 2b). No differences in oxIAA levels were detected among any GFP samples, any ‘no bacteria’ samples, nor any mock AtDAO1 samples (Fig. 2b), suggesting that low levels of non-enzymatic oxidation of IAA occurs in the culture media. Together, these results suggest that incubation with the substrate and the DAO cofactor mixture for 6 hours at 20°C provides the optimal conditions for accumulating enzymatic product. These experimental settings were therefore adopted to develop the method further in later experiments with other enzymes involved in IAA inactivation; only GFP samples were included in the next experiments as a negative control.

**Fig. 3.**
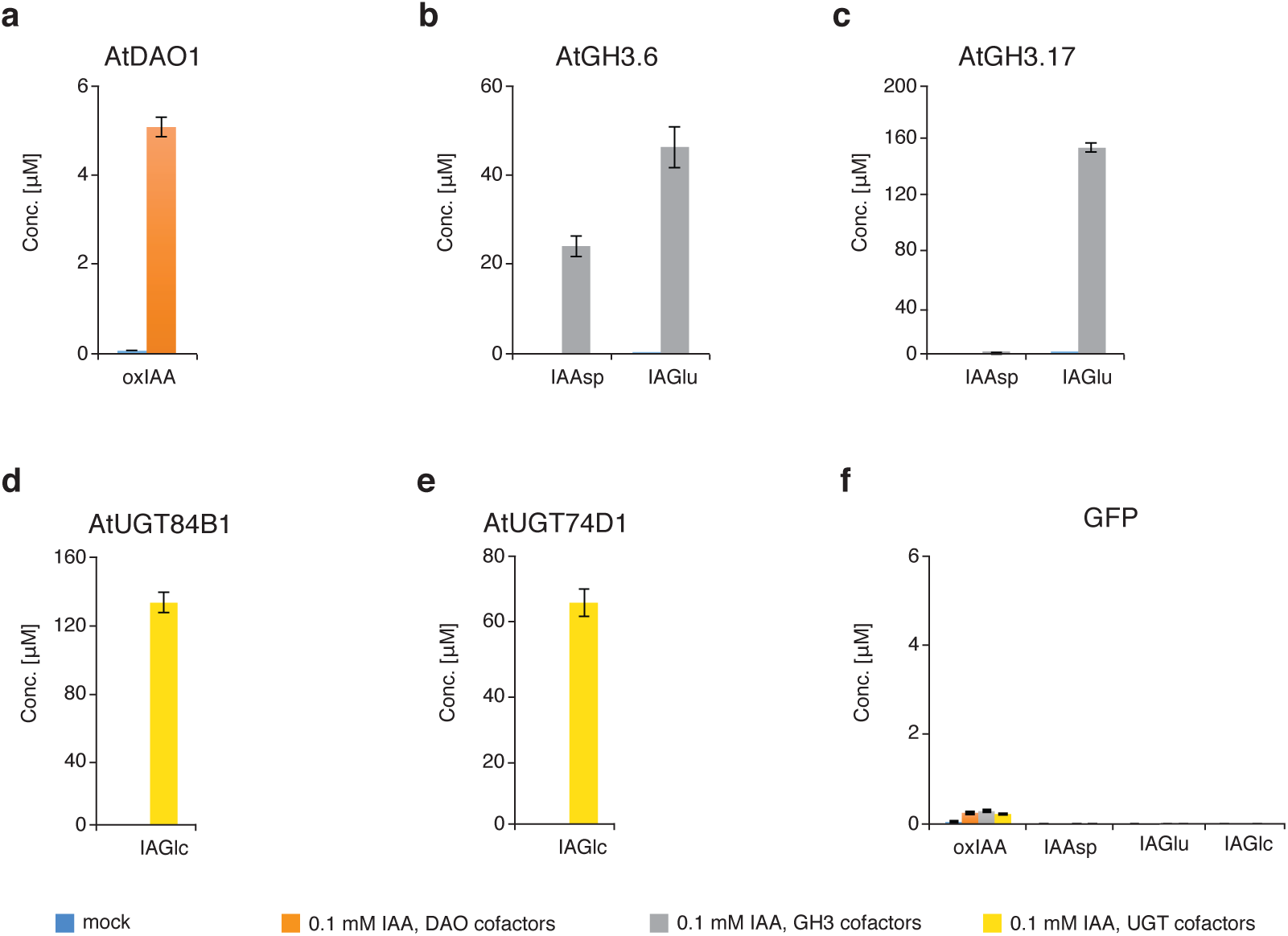
Analysis of IAA metabolites synthesized by recombinant AtDAO1, AtGH3.6, AtGH3.17, AtUGT84B1 and AtUGT74D1 in bacterial assay. **a** oxIAA concentrations in AtDAO1-expressing bacterial cultures. **b, c** IAAsp and IAGlu concentrations in AtGH3.6-(**b**) or AtGH3.17-(**c**) expressing bacterial cultures. **d, e** IAGlc levels measured in AtUGT84B1-(**d**) or AtUGT74D1-(**e**) expressing bacterial cultures. **f** Levels of IAA metabolites in control GFP-expressing bacterial cultures. All bacterial cultures were incubated with 0.1 mM IAA and with the corresponding cofactor mixture for 6 hours at 20°C and only supernatant fractions were analyzed. Bacterial cultures without IAA and cofactor mixture were used as mock samples. Mean ± SD (*n*=3). N.D. = not detectable.

### IAA-conjugating activity of AtGH3.6, AtGH3.17, AtUGT84B1 and AtUGT74D1 in bacterial assay

Earlier work has demonstrated that Group II members of the GH3 family convert IAA into IAA-amino acids [9]. Some of the IAA-amino acid conjugates, such as IAAsp and IAGlu, are the most abundant in Arabidopsis [17] and are believed to be irreversible conjugates that cannot be hydrolyzed to form free IAA [12]. Staswick et al. [9] tested the *in vitro* activity of several recombinant Arabidopsis GH3 proteins and carried out a screening of amino acid preferences using a TLC-based assay by which it was shown that AtGH3.17 prefers Glu over other amino acids, whereas AtGH3.6 exhibits activity with Asp. We therefore examined whether AtGH3.6 and AtGH3.17 could also catalyze the same reaction (Fig. 1b) directly in bacterial cultures that express these recombinant proteins. *AtGH3.6* and *AtGH3.17* genes were expressed in *E. coli* individually and the expression of recombinant fusion proteins was confirmed by Western blot analysis (<u>Additional File 2: Figure S1</u>). Cultures of AtGH3.6- and AtGH3.17-expressing bacteria were tested in a reaction with or without 0.1 mM IAA in combination with the GH3 cofactor mixture (containing as co-substrates, aspartic and glutamic acid, and as cofactors, ATP and MgCl_2_) to determine whether IAAsp and/or IAGlu were formed. Figure 3b shows that the reaction mediated by AtGH3.6 yielded the accumulation of both IAA-amino acid conjugates, IAAsp and IAGlu, which is consistent with previous results reported by Staswick et al. [9]. Interestingly, AtGH3.6 preferred Glu over Asp since IAGlu concentration was higher than IAAsp under our experimental conditions (Fig. 3b). Bacterial cultures expressing AtGH3.17 that were treated with IAA and the GH3 cofactor mixture accumulated very high levels of IAGlu while almost no IAAsp was detected (Fig. 3c), confirming that AtGH3.17 favors Glu over Asp as reported by Staswick et al. [9]. Accumulation of IAAsp and IAGlu was always below the limits of detection in mock samples taken from AtGH3.6- and AtGH3.17-expressing bacterial cultures (Figs. 3b and 3c).

Glucosylation is another important mechanism of IAA conjugation that is implicated in the inactivation of IAA in plants. Members of the UDP-glucosyl transferase (UGT) super family can catalyze glucosylation of plant hormones, including IAA, using UDP-glucose as a co-substrate. Previous studies have demonstrated that two AtUGTs, AtUGT84B1 and AtUGT74D1, can convert IAA to IAGlc *in vitro* (Fig. 1b) [13,14]. To investigate if AtUGT84B1 and AtUGT74D1 could catalyze the glucosylation of IAA in our experimental conditions, we expressed these two genes in *E. coli* individually and confirmed the expression of recombinant fusion proteins by Western blot analysis (<u>Additional File 2: Figure S1</u>). Next, we treated bacterial cultures with or without 0.1 mM IAA in combination with the UGT cofactor mixture (containing as co-substrate, UDPG, and as cofactors, MgSO_4_ and KCl) and measured IAGlc levels. IAGlc accumulated in samples taken from cultures of bacteria transformed with either *AtUGT84B1* or *AtUGT74D1* that were treated with 0.1 mM IAA and UGT cofactor mixture, while IAGlc levels were below the limits of detection in samples taken from the same bacterial cultures that were not treated with IAA and UGT cofactor mixture (Figs. 3d and 3e). IAAsp, IAGlu and IAGlc levels were below the limits of detection in all GFP samples treated with IAA and cofactor mixtures, and in all mock GFP samples (Fig. 3f). Taken together, these results demonstrate that AtGH3 and AtUGT proteins heterologously expressed in *E. coli* cells can efficiently conjugate IAA with amino acids and sugar, respectively, by performing the enzymatic assay with IAA and specific cofactor mixtures directly in bacterial cultures.

### OxIAA-conjugating activity of AtUGT84B1 and AtUGT74D1 and competition assay

Conjugation of oxIAA with glucose to form oxIAGlc is an additional and relevant metabolic step in the oxIAA pathway. oxIAGlc, together with oxIAA, are the major degradation products of IAA in Arabidopsis [12,17,18], suggesting that this reaction also contributes to IAA homeostasis. Previous studies have demonstrated that recombinant AtUGT74D1 protein converts not only IAA to IAGlc [14], but also oxIAA to oxIAGlc (Fig. 1b) [15]. To verify whether recombinant AtUGT proteins exhibit glucosyl transferase activity towards oxIAA under our experimental conditions, we treated AtUGT84B1- and AtUGT74D1-expressing bacterial cultures with 0.1 mM oxIAA in combination with the UGT cofactor mixture. oxIAGlc was detected in cultures that expressed AtUGT84B1 and AtUGT74D1, after treatment with oxIAA (Figs. 4a and 4b), indicating that these two AtUGTs can catalyze the glucosylation of oxIAA. To determine the substrate preference of these two AtUGTs, we performed a competition experiment in which cultures of AtUGT84B1- and AtUGT74D1-expressing bacteria were treated simultaneously with the same concentration of IAA and oxIAA and with the UGT cofactor mixture. When IAA and oxIAA competed with each other, IAA was found to be a better substrate as IAGlc levels were much higher than oxIAGlc in samples from both bacterial cultures expressing either of the two AtUGTs (Figs. 4a and 4b). IAGlc and oxIAGlc were always below the limits of detection in GFP samples as well as in both AtUGTs mock samples (Fig. 4). Our finding is in contrast to that reported by Tanaka et al. [15]; they showed that recombinant AtUGT74D1 had a higher specificity towards oxIAA than IAA *in vitro*. The discrepancy between these findings could be due to the different experimental conditions that were adopted to study the enzymatic activity.

**Fig. 4.**
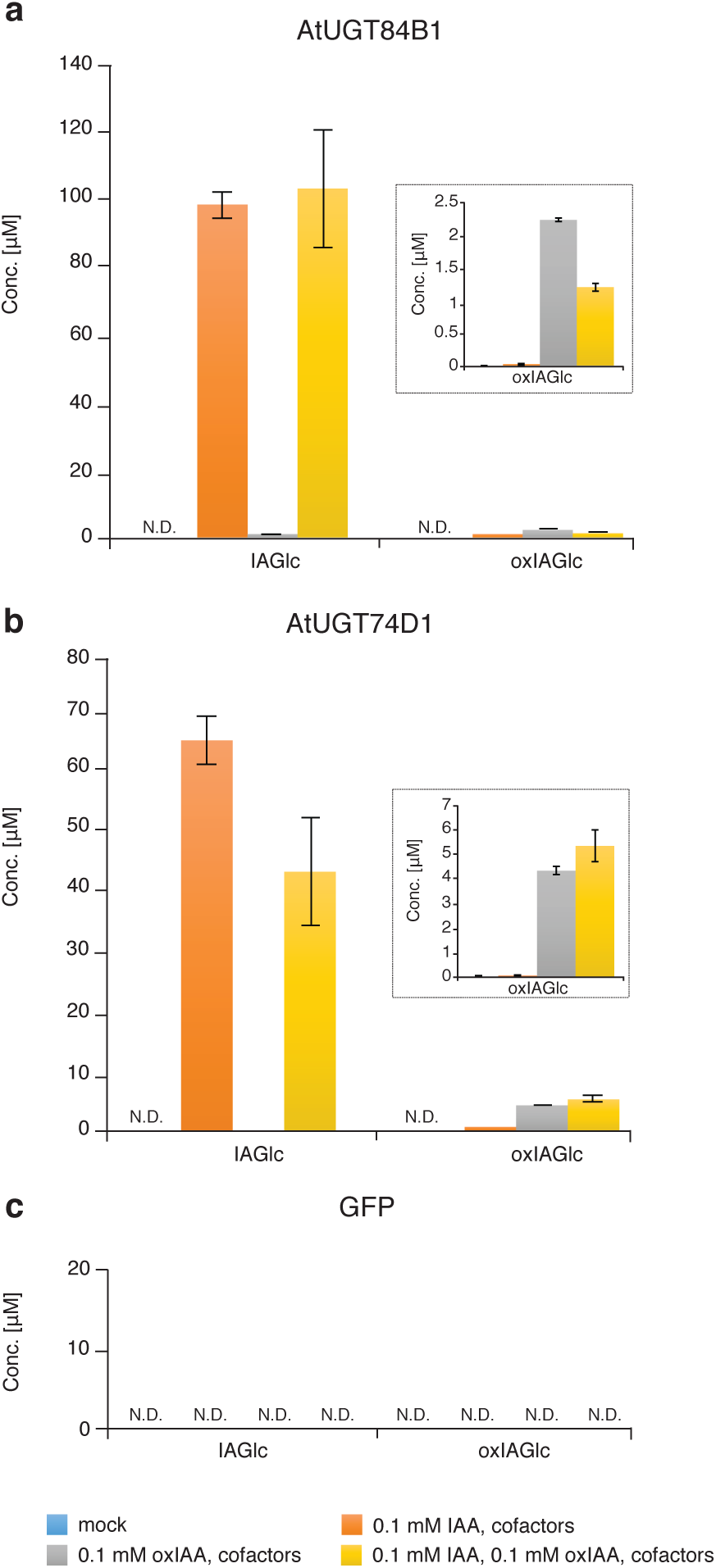
Analysis of IAGlc and oxIAGlc synthesized by AtUGTs in a competition assay. **a, b** IAGlc and oxIAGlc (inset) concentrations measured in supernatant fractions from AtUGT84B1-(**a**) or AtUGT74D1-(**b**) expressing bacterial cultures. **c** Levels of IAGlc and oxIAGlc in control GFP-expressing bacterial cultures. All the bacterial cultures were incubated with 0.1 mM IAA and/or oxIAA and with UGT cofactor mixture for 6 hours at 20°C and only supernatant fractions were analyzed. Bacterial cultures without IAA and UGT cofactor mixture were used as mock samples. Mean ± SD (*n*=3). N.D. = not detectable.

Taken together, these results provide evidence that our method is suitable for testing different substrates by supplementing the compounds of interest, individually or simultaneously, directly in the bacterial cultures expressing the recombinant protein. In the present study, we were also able to show that AtUGT84B1 can catalyze not only the glucosylation of IAA but also of oxIAA to their corresponding glucosides. To the best of our knowledge, this is the first report in which AtUGT84B1 glucosyl transferase activity has been tested and demonstrated toward oxIAA.

## Conclusions

The development of a rapid alternative method for screening heterologously expressed Arabidopsis enzymes involved in IAA inactivation is described. Our method takes advantage of the fact that recombinant Arabidopsis proteins that inactivate IAA retain their biological activity when expressed in bacteria. Our results show that an enzymatic assay can be conducted directly in the bacterial cultures expressing recombinant proteins because the expected IAA metabolites are rapidly and abundantly produced upon supplementation of substrate and specific cofactor mixtures to engineered bacterial cultures. IAA metabolites that accumulate in culture media are identified and quantified using UHPLC-ESI-MS/MS by adding corresponding internal standards. Prior *in vitro* methods for assaying protein activity are based on expressing Arabidopsis enzymes in a recombinant form in bacteria or yeast followed by recombinant protein purification. However, the extraction and purification of recombinant proteins is a laborious and expensive process and a major obstacle when assessing *in vitro* assays. The enzymatic method described here eliminates this technical constraint and represents a flexible system that is suitable for rapidly screening the activity of different hormone-modifying enzymes. It should be noted that, because it is impossible to quantify recombinant proteins directly in bacteria, at this stage our method cannot guarantee a precise kinetics of the enzymes. Instead, the method proposed here provides an easy but powerful alternative to screen any putative orthologous genes from non-model species, allowing the characterization of genes from the newly sequenced genomes. As such it could be used to test for enzyme affinity toward different substrates, such as natural and synthetic molecules. However, due to its versatility, our method could also be adopted for testing specific enzyme inhibitors, such as compounds that are of interest in agriculture and horticulture.

## Supporting information

Supplemental Figue 1

## Additional material

**Additional File 1: Table S1.** List of primers used in this work for cloning.

**Additional File 2: Figure S1.** Western blot analysis of IPTG-induced *E. coli* harboring recombinant AtDAO1, AtGH3.6, AtGH3.17, AtUGT84B1, AtUGT74D1 or GFP construct. Bacterial cultures were treated with 0.1 mM IPTG and incubated 6 hours either at 20°C. For bacterial enzymatic assays, recombinant protein expression was obtained with over-night incubation at 20°C in the presence of 0.1 mM IPGT. Calculated MW of recombinant proteins: 35.7 kDa (AtDAO1), 69.9 kDa (AtGH3.6 and AtGH3.17), 51.7 kDa (AtUGT84B1), 51.1 kDa (AtUGT74D1) and 27 kDa (GFP).

**Additional File 3: Table S2.** List of targeted compounds with their respective retention times (RT) and MRM transition. Indole-3-acetic acid (IAA), oxindole-3-acetic acid (oxIAA), indole-3-acetyl-L-aspartic acid (IAAsp), indole-3-acetyl glutamic acid (IAGlu), indole-3-acetyl-1-glucosyl ester (IAGlc) and oxindole-3-acetyl-1-glucosyl ester (oxIAGlc).

**Additional File 4: Figure S2.** Analysis of IAA and oxIAA background concentrations in liquid media containing AtDAO1- or GFP-expressing bacteria or no bacteria prior supplementation with exogenous IAA (T0). Mean ± SD (*n*=3).

**Additional File 5: Dataset S1.** Raw data from mass spectrometry analysis.

## Abbreviations

CCD: charge-coupled device
DAO: DIOXIGENASE OF AUXIN OXIDATION
ECL: enhanced chemiluminiscent
GH3: GRETCHEN HAGEN 3
HRP: horseradish peroxidase
IAA: indole-3-acetic acid
IAGlc: indole-3-acetyl-1-glucosyl ester
IAGlu: indole-3-acetyl-L-glutamic acid
IAAsp: indole-3-acetyl-L-aspartic acid
IPTG: isopropyl-β-D-thiogalactopyranoside
oxIAA: oxindole-3-acetic acid
oxIAGlc: oxindole-3-acetyl-1-glucosyl ester
TLC: thin-layer chromatography
Trp: tryptophan
UGT: UDP glucosyl transferase
UHPLC-ESI-MS/MS: ultrahigh-performance liquid chromatography coupled to electrospray ionization tandem spectrometry.

## Acknowledgements

This research was supported by grants from the Swedish research councils FORMAS, VR, Kempestiftelserna, and the Knut and Alice Wallenberg Foundation (to KL and CB). The authors would like to thank the Swedish Metabolomics Center (SMC, Umeå, Sweden) for access to instrumentation.

## Authors’ contributions

FB and SC designed the experiments. FB, SC and JS performed all the experiments. FB and SC wrote the article with the input from all authors. All authors read and approved the final article for publication.

## References

1. Vanneste S, Friml J. Auxin: A Trigger for Change in Plant Development. Cell. 2009;136(6):1005–16.

2. Friml J, Vieten A, Sauer M, Weijers D, Schwarz H, Hamann T, et al. Efflux dependent auxin gradients establish the apical-basal axis of Arabidopsis. Nature. 2003;426(6963):147–53.

3. Benková E, Ivanchenko MG, Friml J, Shishkova S, Dubrovsky JG. A morphogenetic trigger: is there an emerging concept in plant developmental biology? Trends Plant Sci. 2009;14(4):189–93.

4. Petersson S V, Johansson AI, Kowalczyk M, Makoveychuk A, Wang JY, Moritz T, et al. An auxin gradient and maximum in the Arabidopsis root apex shown by high-resolution cell-specific analysis of IAA distribution and synthesis. Plant Cell. 2009;21(6):1659–68.

5. Ljung K. Auxin metabolism and homeostasis during plant development. Development. 2013;140(5):943–50.

6. Zhao Y. Essential Roles of Local Auxin Biosynthesis in Plant Development and in Adaptation to Environmental Changes. Annu Rev Plant Biol. 2018;69(1).

7. Ludwig-Müller J. Auxin conjugates: Their role for plant development and in the evolution of land plants. J Exp Bot. 2011;62(6):1757–73.

8. Zhang J, Peer WA. Auxin homeostasis: The DAO of catabolism. J Exp Bot. 2017;68(12):3145–54.

9. Staswick PE, Serban B, Rowe M, Tiryaki I, Maldonado MT, et al. Characterization of an Arabidopsis enzyme family that conjugates amino acids to indole-3-acetic acid. Plant Cell. 2005;17:617–627.

10. Westfall CS, Herrmann J, Chen Q, Wang S, Jez JM. Modulating plant hormones by enzyme action. Plant Signal Behav. 2011;5(12):1607–12.

11. Ruiz Rosquete M, Barbez E, Kleine-Vehn J. Cellular auxin homeostasis: Gatekeeping is housekeeping. Mol Plant. 2012;5(4):772–86.

12. Ostin A, Kowalyczk M, Bhalerao RP, Sandberg G. Metabolism of indole-3-acetic acid in Arabidopsis. Plant Physiol. 1998;118(1):285–96.

13. Jackson RG, Lim EK, Li Y, Kowalczyk M, Sandberg G, Hoggett J, et al. Identification and biochemical characterization of an Arabidopsis indole-3-acetic acid glucosyltransferase. J Biol Chem. 2001; 276:4350–4356.

14. Jin SH, Ma XM, Han P, Wang B, Sun YG, Zhang GZ, et al. UGT74D1 Is a Novel Auxin Glycosyltransferase from Arabidopsis thaliana. PLoS One. 2013;8(4):1–11.

15. Tanaka K, Hayashi KI, Natsume M, Kamiya Y, Sakakibara H, Kawaide H, et al. UGT74D1 catalyzes the glucosylation of 2-oxindole-3-acetic acid in the auxin metabolic pathway in arabidopsis. Plant Cell Physiol. 2014;55(1):218–28.

16. Kowalyczk M, Sandberg G. Quantitative analysis of indole-3-acetic acid metabolites in Arabidopsis. Plant Physiol. 2001;127:1845–1853.

17. Novák O, Hényková E, Sairanen I, Kowalczyk M, Pospíšil T. Tissue-specific profiling of the Arabidopsis thaliana auxin metabolome. Plant J. 2012;72:523–536.

18. Porco S, Pěnčík A, Rashed A, Voß U, Casanova-Sáez R, Bishopp A, et al. Dioxygenase-encoding AtDAO1 gene controls IAA oxidation and homeostasis in Arabidopsis. Proc Natl Acad Sci. 2016;113(39):11016–21.

19. Zhang J, Lin JE, Harris C, Campos Mastrotti Pereira F, Wu F, Blakeslee JJ, et al. DAO1 catalyzes temporal and tissue-specific oxidative inactivation of auxin in Arabidopsis thaliana. Proc Natl Acad Sci. 2016;113(39):11010–5.

20. Peer WA, Cheng Y, Angus M. Evidence of oxidative attenuation of auxin signalling. J Exp Bot. 2013;64:2629–2639.

21. Pencik A, Simonovik B, Petersson S V., Henykova E, Simon S, Greenham K, et al. Regulation of Auxin Homeostasis and Gradients in Arabidopsis Roots through the Formation of the Indole-3-Acetic Acid Catabolite 2-Oxindole-3-Acetic Acid. Plant Cell. 2013;25(10):3858–70.

22. Stepanova AN, Alonso JM. Auxin catabolism unplugged: Role of IAA oxidation in auxin homeostasis. Proc Natl Acad Sci. 2016;113(39):10742–4.

23. Jackson RG, Kowalczyk M, Li Y, Higgins G, Ross J, Sandberg G, et al. Over-expression of an Arabidopsis gene encoding a glucosyltransferase of indole-3-acetic acid: Phenotypic characterisation of transgenic lines. Plant J. 2002;32(4):573–83.

24. Böttcher C, Keyzers RA, Boss PK, Davies C. Sequestration of auxin by the indole-3-acetic acid-amido synthetase GH3-1 in grape berry (Vitis vinifera L.) and the proposed role of auxin conjugation during ripening. J Exp Bot. 2010;61(13):3615–25.

25. Ludwig-Müller J, Jülke S, Bierfreund NM, Decker EL, Reski R. Moss (Physcomitrella patens) GH3 proteins act in auxin homeostasis. New Phytol. 2009;181(2):323–38.

26. Chen Q, Zhang B, Hicks LM, Wang S, Jez JM. A liquid chromatography-tandem mass spectrometry-based assay for indole-3-acetic acid-amido synthetase. Anal Biochem. 2009;390(2):149–54.

27. Chen Q, Westfall CS, Hicks LM, Wang S, Jez JM. Kinetic Basis for the Conjugation of Auxin by a GH3 Family. J Biol Chem. 2010;285(39):29780–6.

28. Zhang S-W, Li C-H, Cao J, Zhang Y-C, Zhang S-Q, Xia Y-F, et al. Altered Architecture and Enhanced Drought Tolerance in Rice via the Down-Regulation of Indole-3-Acetic Acid by TLD1/OsGH3.13 Activation. Plant Physiol. 2009;151(4):1889–901.

29. Drábková LZ, Dobrev PI, Motyka V. Phytohormone profiling across the bryophytes. PLoS One. 2015;10(5):1–19.

30. Žižková E, Kubeš M, Dobrev PI, Přibyl P, Šimura J, Zahajská L, et al. Control of cytokinin and auxin homeostasis in cyanobacteria and algae. Ann Bot. 2017;119(1):151–66.

31. Pencík A, Casanova-Sàez R, Pilarovà V, Žukauskaite A, Pinto R, et al. Ultra-rapid auxin metabolite profiling for high-throughput mutant screening in Arabidopsis. J Exp B. 2018;69(10):2569–2579.

32. Zhao Z, Zhang Y, Liu X, Zhang X, Liu S, et al. A role for a dioxigenase in auxin metabolism and reproductive development in rice. Dev Cell. 2013;27:113–122.

