## Supplemental Figue 1 for "A bacterial assay for rapid screening of IAA catabolic enzymes"

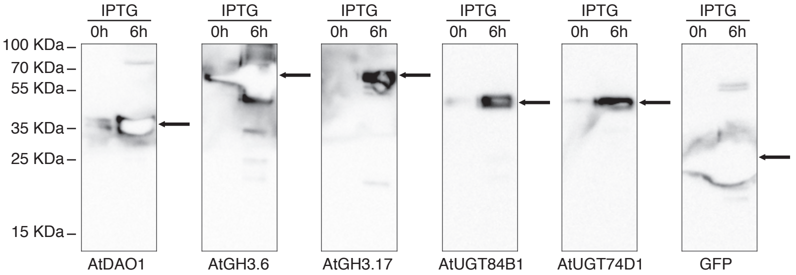


**Figure S1.** Western blot analysis of IPTG-induced *E. coli* harboring recombinant AtDAO1, AtGH3.6, AtGH3.17, AtUGT84B1, AtUGT74D1 or GFP construct. Bacterial cultures were treated with 0.1 mM IPTG and incubated 6 hours at 20°C. For bacterial enzymatic assays, recombinant protein expression was obtained with over night incubation with IPTG at 20°C in the presence of 0.1 mM IPTG. Calculated MW of recombinant proteins: 35.7 kDa (AtDAO1), 69.9 kDa (AtGH3.6 and AtGH3.17), 51.7 kDa (AtUGT84B1), 51.1 kDa (AtUGT74D1) and 27 kDa (GFP).
