## Supplemental Figure 2 for "A bacterial assay for rapid screening of IAA catabolic enzymes"

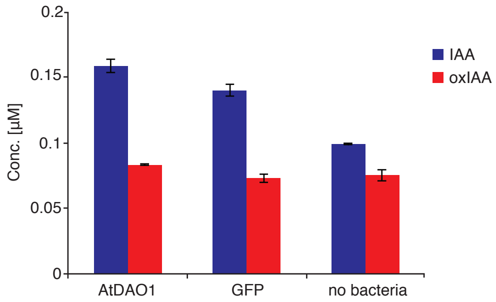


**Figure S2.** Analysis of IAA and oxIAA background concentrations in liquid media containing AtDAO1- or GFP-expressing bacteria or no bacteria prior supplementation with exogenous IAA (T0). Mean ± SD (*n*=3).
